# Representational pattern similarity of electrical brain activity reveals rapid and specific prediction during language comprehension

**DOI:** 10.1101/2020.04.23.058552

**Authors:** Ryan J. Hubbard, Kara D. Federmeier

## Abstract

Predicting upcoming stimuli and events is a critical function of the brain, and understanding the mechanisms of prediction has thus become a central topic in neuroscientific research. Language provides a fertile testing ground for examining predictive mechanisms, as comprehenders use context to predict different features of upcoming words. Although there is a substantive body of research on prediction in language, many aspects of the mechanisms of prediction remain elusive, in part due to a lack of methodological tools to probe prediction formation in the moment. To elucidate what features are neurally pre-activated and when, we used representational similarity analysis (RSA) on data from a sentence reading task (Federmeier et al., 2007). We compared EEG activity patterns elicited by expected and unexpected sentence final words to patterns from the preceding words of the sentence, in both strongly and weakly constraining sentences. Pattern similarity with the final word was increased in an early time window (suggestive of visual feature activation) following the presentation of the pre-final word, and this increase was modulated by both expectancy and constraint (greatest for strongly constrained expected words). This was not seen at earlier words, suggesting that predictions are precisely timed. Additionally, pre-final word activity – the predicted representation - had negative similarity with later final word activity, but only for strongly expected words. Together, these findings shed light on the mechanisms of prediction in the brain: features of upcoming stimuli are rapidly pre-activated following related cues, but the predicted information may receive reduced subsequent processing upon confirmation.

## Introduction

Theories of cognition and neural functioning increasingly build in an important role for anticipatory processing – i.e., prediction. Indeed, some have postulated that a core mechanism of neural coding involves higher-level cortical systems in the brain attempting to predict and explain input at lower levels in a hierarchical fashion (*predictive coding* [1–3]). One area that has proven to be a particularly rich testing ground for understanding the import – and limitations – of predictive processing is language comprehension. When listening to or reading language, people can use contextual cues and prior knowledge to generate predictions about upcoming words in order to support rapid and efficient comprehension and communication [4–7]. The contents of these predictions can be multifaceted in nature, including orthographic [8–9], phonological [10–11], semantic [12–13], and morphosyntactic [14–15] features of words. However, such anticipatory processes are not always engaged [16–17], suggesting that the brain flexibly allocates resources to predict information to the extent that the environment allows it and as a function of the utility of those predictions for the task at hand.

Studies of prediction in language using behavioral [18–20] and eyetracking [21–23] measures combine with a sizeable literature that has focused on neural responses to predictable and unpredictable words as measured via electroencephalography (EEG), including event-related potentials (ERPs [17, 24–26]), or magnetoencephalography (MEG [14, 27–29]). This work has established that both the constraint of a context (i.e., how much it narrows expectations and allows a strong, consistent prediction) and the probability of the encountered word in its context modulate brain responses [27, 30–31]. Effects can even be seen on determiners or modifiers, when these have specific gender or phonological characteristics (e.g. “*a* kite”) that are consistent or inconsistent with an anticipated noun (“*an* … kite”) [10, 15, 32–35], as well as within 150 ms following word onsets that can differentiate between words with many possible continuations and words with few [36–37]. However, in much extant work prediction has been inferred from brain responses that obtain after a more or less predictable word (or a determiner/modifier carrying those features) is encountered, making it difficult to know precisely when and how predictions were formed.

To try to instead capture processing at the time that predictions are being made, some work has examined event-related activity differences elicited by a verb or adverb, as a function of whether it does or does not afford a prediction for a target, sentence-final word [38–39]. These studies report more negative N400s for more predictive preceding words, suggesting that predictability can affect processing in advance; however, it is unclear if these responses do or not reflect pre-activation of specific features of upcoming information. Other studies have employed novel paradigm manipulations in order to examine anticipatory processing. For instance, León-Cabrera, Rodríguez-Fornells, & Morís [40] presented participants with sentences that varied in contextual constraint. By imposing a 1000 ms delay before the target word, they were able to observe slow negative potentials that were sensitive to constraint. Dikker & Pylkkänen [41] implemented a picture-noun matching task to examine pre-activation of lexical features, in which more or less predictive pictures preceded related nouns; they found MEG activity differences based on predictability 400 ms prior to noun onset. However, it is unclear if the effects in these studies arise due to the unique demands of the task, and thus if the same processes would be observed in more natural language comprehension settings.

Examining ERP and MEG responses to prior words or time windows in isolation presents difficulties in separating predictive processing of the upcoming information from reactive processing of the preceding information, or more general effects of constraint. An optimal method for examining pre-activation would consider both neural activity prior to and following the target stimulus to determine if there is similarity in neural processing, and if that similarity varies with predictability. If the brain pre-activates features of upcoming words, then patterns of neural activity specifically related to processing that word should be present in advance, and comparing these patterns should reveal similarity graded by constraint and match to expectation. Representational similarity analysis (RSA) presents a promising solution [42–44]. With this technique, multivariate patterns of neural activity are compared with a correlational approach, which can be performed across the neural time-series or across electrodes. This method allows not only for identification of both temporal and spatial patterns of similarity, but also for detection of more subtle but statistically separable neural states than may be found with more conventional ERP analyses [45].

A recent study used RSA of MEG data to investigate pre-activation of semantic features in a language comprehension paradigm [29]. Specifically, participants read strongly constraining sentences that were constructed in pairs, such that within-pair sentences predicted the same sentence-final critical word (e.g. “In the crib there is a sleeping…” and “In the hospital there is a newborn…” both predict the word “baby”) and between-pair sentences did not. A greater increase in neural similarity was found for within-pair sentences, suggesting greater similarity of neural patterns across sentences wherein the same word was predicted. However, it is unclear whether this was entirely due to prediction, or at least partly reflected that the brain was in a more similar state due to the shared semantics of within-pair sentences. Additionally, even “pseudo-repetitions” of words that were expected but never presented can lead to a lingering representation in the brain, despite intervening sentences [46], which may have influenced pattern similarity.

In this paper, we target prediction specifically by comparing pre-final word activity with post-final word activity in sentences that varied in constraint and had expected or unexpected endings. We thus circumvent the issue of semantic similarity across sentences that yield similar predictions, because, in this case, the pattern comparisons for expected and unexpected words are within the same sentence context. If the input of the pre-final word cues pre-activation of features of the final word, then some aspects of the neural representation of the final word should appear during the processing of the pre-final word, which will be detected with RSA. Moreover, critically, if this correlation is, indeed, due to prediction-related activations, then similarity for expected words should be greater when sentential constraint is higher, and similarity should be reduced or potentially abolished when the ending is unexpected, as the neural representation of the final word may no longer match with the pre-activated representation. Thus, the effect of predictability can be tested by examining graded similarity across levels of constraint, and the effect of disconfirming predictions can be tested by comparing similarity for expected and unexpected outcomes. Finally, the timing of prediction generation can be examined by assessing the similarity of final word activity patterns with patterns elicited by words preceding the pre-final word.

We also employ a time generalization analysis in order to examine the time-course or development of predictions over time [47–48]. Pattern similarity may increase gradually across time as the onset of the target word approaches, or may come on and offline more rapidly [29, 41]. Additionally, this analysis method allows us to probe the fate of predicted representations after encountering the predicted word itself. Representations of words that were previously predicted have been found, downstream, to be impoverished compared to unpredicted words, suggesting later processing of word representations differs based on predictability [49–50]. This difference in processing may be observable by analyzing representational similarity of pre-final word activity with later time windows of post-final word activity.

## Methods

This paper uses novel techniques to reanalyze data from the study reported in Federmeier et al. (2007). Further specifics of the methodology can be found in that publication.

### Participants

32 right-handed individuals participated in the experiment in exchange for course credit or cash. One individual was dropped due to technical issues with importing the data, resulting in a total of 31 participants in the analysis. All participants reported normal or corrected vision and had no history of any neurological or psychiatric disorder. Mean age was 20 years (range 18-28 years), and 16 of the participants were female. The study was approved by the local ethics committee, and all participants provided written informed consent and were debriefed following participation.

### Design and Procedure

Participants read 282 sentences that varied in contextual constraint (for bin-based analyses, divided into high constraint, cloze > 0.67, and low constraint, cloze < 0.42), and ended with either an expected or unexpected (cloze ≈ 0.03), but plausible word. Stimuli were counter-balanced into two lists, such that half of the sentences completed by an expected ending in one list were completed by an unexpected ending in the second list, and vice versa. Thus, there were four conditions of sentence final words: strong constraint expected (SCE), strong constraint unexpected (SCU), weak constraint expected (WCE), and weak constraint unexpected (WCU), with approximately 70 sentences in each condition. Sentence frames were matched in length, and lexical properties (word length, word frequency) of sentence endings were matched.

Words prior to the final word of the sentence, or pre-final words, were primarily made up of determiners or prepositions (“the”, “a”, “his”, etc; 65% of pre-final stimuli). Pre-final words did not reliably differ in length across sentence constraint (*p* = 0.06), but did differ in log frequency (*p* < 0.01). Additional linear mixed effect model analyses were run to include lexical confounds as predictors for experimental effects of interest. Mixed effect models were conducted in R, using the lme4 package [51], and statistical significance of fixed effects were estimated with t-tests using the Satterthwaite method in the lmerTest package [52].

Association strength between words within sentences and sentence-ending words was measured using the Edinburgh Associative Thesaurus. For each sentence in each condition, the forward and backward association was calculated between each word in the sentence and the sentence final word, and the number of instances of associations greater than 0.2 were counted. Overall, there were few associations between sentence words and sentence-ending words: 4% of SCE sentences contained at least one association greater than 0.2, and for all other conditions (SCU, WCE, WCU) only 1% of sentences contained at least one association. The mean association strength between sentence final words and all words in the sentence was less than 0.005 for each of the four conditions. Thus, observed effects are unlikely to arise simply due to word level associations.

Participants viewed the sentences on a 21″ CRT monitor in an electrically shielded booth. Each sentence was presented word-by-word in the center of the screen, with each word appearing for 200 ms with an interstimulus interval of 300 ms. Sentences were separated by a 3 s pause. Participants were instructed to attend to and read the sentences for comprehension, and that they would be asked questions about what they had read at the end of the experiment.

### EEG Recording and Processing

EEG data was recorded from 26 tin electrodes embedded into a flexible elastic cap distributed over the scalp in an equidistant arrangement. Additional electrodes included one on each mastoid, one on each outer canthus of the eye (for monitoring eye movements), and one below the lower eyelid of the left eye (for monitoring blinks). Electrode impedances were kept below 5 kΩ. Signals were amplified by a Grass amplifier with a bandpass filter of 0.01-100 Hz, and a sampling rate of 250 Hz. During recording, the left mastoid electrode was used as a reference; offline, the data were re-referenced to the average of the left and right mastoid electrodes.

Raw EEG time series were filtered with a 0.2-60 Hz digital Butterworth bandpass filter with a 12 dB/oct roll-off. Filter parameters were chosen *a priori* to remove high frequency noise and large drifts but retain some higher frequencies that could potentially contribute meaningful variance to the RSA; however, a second analysis with a 0.2-30 Hz filter produced nearly identical results. Note that the high-pass filter was implemented to reduce noise from low frequency drifts but was not so high as to induce confounds in the similarity analysis. Namely, the time windows used in this analysis were similar to those used in the ERP analysis, and previous work has demonstrated that a 0.2 high pass filter does not produce distortions in electrophysiological measurements over these time scales [53]. To correct ocular artifacts, the data were decomposed into independent components with the AMICA algorithm [54]. Component time-courses that correlated with a bipolar vertical electrooculogram (EOG) channel at Pearson *r* > 0.6 were removed, and the data were reconstructed with the remaining components. The corrected data were then submitted to a sliding window artifact scan to identify extreme amplitude excursions (>90 μV). Any trial in which either the pre-final word or final word was marked as an artifact was excluded from analysis. RSA results could potentially be influenced by differences in trial numbers [55], and so for each participant the number of trials in each condition was equated by randomly dropping trials from conditions with more trials than the condition with the minimum number. This resulted in an average of 64 trials in each condition for each participant.

The remaining trials were corrected with a z-scoring baseline correction method [56], in which pre-trial baseline periods are fused together and used to z-score the trial data. This method reduces potential biases of single-trial normalization techniques [57]. Separate baselines were created for strong constraint and weak constraint sentences to reduce any contamination from sentential constraint (e.g. strong constraint baselines may differ somewhat from weak constraint baselines).

Additionally, both the pre-final word and final word data were corrected with pre-final word baseline data, so as not to introduce any bias by using pre-final word data both as a baseline and in the similarity analysis.

### Spatial Representational Similarity Analysis

Spatial RSA is focused on the similarity of neural activity across the scalp at each timepoint of two time series. For each trial of data from each participant, the vector of amplitudes from each of the 26 scalp channels of the pre-final word data was correlated (Pearson’s *r*) with the vector of channel data of the final word data at each and every time-point from 1 to 500 ms post-word onset. This resulted in a time-series of correlations between pre-final and final word activity for each trial. For bin-based analyses, these time-series were then averaged across trials within each of the four conditions (SCE, SCU, WCE, and WCU) for each participant, and grand averages were created by averaging across participants. Additionally, a grand average across all four conditions was created, which was used to identify timepoints of interest for analysis so as not to bias our decision by viewing the condition data [58]. While this method is subjective, it is unbiased in terms of condition, and allows for greater statistical power than the more conservative mass univariate approaches. Peak similarity in the grand average was observed at 185 ms; statistical analyses were then conducted in a 50 ms window around that peak, using repeated-measures ANOVAs for testing factors of expectancy and constraint.

### Time Generalization Analysis

Time generalization follows the same steps as the spatial RSA, but the channel activity at each time-point of the pre-final word is correlated with the channel activity of every time-point of the final word, producing a matrix of correlations with the matching time-points on the diagonal. Here, only the first 300 ms of the pre-final word activity was correlated with 1-500 ms of the final word activity. The later time-points of the pre-final word are close in proximity to the early time-points of the final word, and thus the overall similarity is greatly increased; however, this is likely not due to prediction, and the large correlation values produced could influence the results of statistical analyses. Thus, the time-range of the pre-final word was limited to 1 to 300 ms to avoid this issue.

Time-generalization matrices were created for each trial and averaged across trials for each of the four conditions for each participant. The resultant average matrices were submitted to cluster-based permutation analyses [59] to test for significant differences in similarity between two conditions. Here, *t*-tests were performed at each time-point testing for differences between conditions. Clusters were identified in the time x time matrix by grouping adjacent time-points where the *t*-test was significant (*p* < 0.05), and the magnitude of each observed cluster was determined by summing the *t*-values within the cluster. A surrogate distribution was then created by shuffling the subject labels, performing *t-*tests at each time-point, identifying significant clusters, and recording the largest cluster statistic. The largest statistic was recorded as both a positive and negative value in order to perform a two-sided test (the null hypothesis distribution was assumed to be symmetric). This shuffling procedure was repeated 1000 times, and the observed clusters of the actual data were then compared to the surrogate distribution of cluster statistics to test for significance. The observed clusters were considered significant if 97.5% of the surrogate cluster values were smaller than the observed cluster value, or if 97.5% of the surrogate cluster values were larger than the observed cluster value. Multiple permutation tests were performed in order to examine differences between the four conditions.

### Temporal Representational Similarity Analysis

Temporal RSA is focused on the similarity of two neural time series at each channel across the scalp, allowing for visualization of the topography of the similarity. For each trial of data from each participant, the vector of amplitudes from the 75 ms time window (150-225 ms) following the pre-final word was correlated with the 150-225 ms time series following the final word at each of the 26 scalp channels. Note that the window used for the temporal RSA was slightly larger than that used for the spatial RSA. The window was widened in order to include more points in the temporal RSA correlation for a more stable estimate. This analysis resulted in a scalp map of correlations between pre-final and final word activity for each trial. As with the spatial RSA, these scalp maps were then averaged across trials within each of the four conditions for each participant and averaged across participants to create a grand average scalp map.

This method was additionally extended to a sliding window approach in order to explore the results from the spatial generalization analysis. For the correlation approach to be possible, the correlated time series must be the same length; thus, we used 60 ms windows of time from both the pre-final word and the final word activity. A 60 ms time window (slightly larger than the original window used for spatial RSA) was used to capture the entire extent of the significant clusters observed. The time series of pre-final word activity correlated with 60 ms windows of post-final word activity in successive 4 ms steps for each trial. The scalp maps at each time step were then averaged across trials within each of the four conditions.

## Results

EEG was recorded while participants read sentences that varied in contextual constraint and ended with either an expected or unexpected word (Fig 1A). We used spatial RSA to compare neural response patterns to pre-final words and final words by correlating the amplitude values across sensors at each timepoint of the two time-series and averaging the resulting similarity time-series within each condition (Fig 1B).

**Figure 1.**
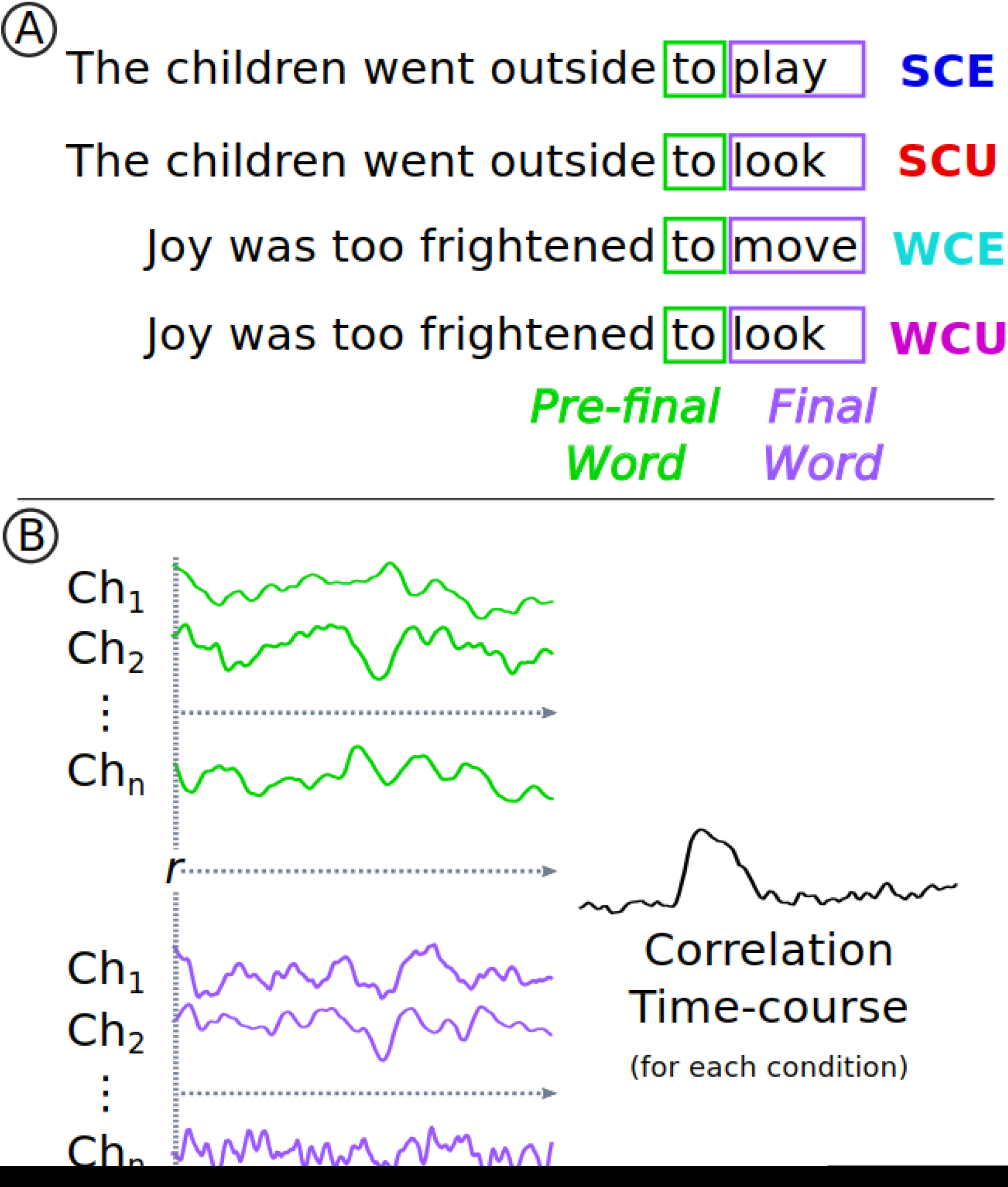
Example of experimental materials and RSA diagram. **A)** Examples of sentences from each of the four conditions: Strong Constraint Expected (SCE), Strong Constraint Unexpected (SCU), Weak Constraint Expected (WCE), and Weak Constraint Unexpected (WCU). The sentence final words are highlighted in purple, and the pre-final words are highlighted in green. **B)** In spatial RSA, the vector of EEG channel activity at the first time-point of the pre-final word (shown in green) is correlated with the vector of activity at the first time-point of the final word (shown in purple). EEG activity correlations are calculated at each successive time-point, resulting in a time-course of correlations.

### Spatial RSA

Spatial RSA revealed a peak in neural similarity beginning around 100 ms and continuing to about 350 ms following word onset (Fig 2A). A repeated measures ANOVA on average similarity values between 160-210 ms (50 ms around the grand average peak) with factors of constraint (high and low) and expectancy (expected and unexpected) found significant main effects of constraint (F_(1,30)_ = 12.32, *p* < 0.01) and expectancy (F_(1,30)_ = 7.67, *p* < 0.01), with no significant interaction (F_(1,30)_ = 0.45, *p* = 0.48). The largest effect was the increase in similarity for expected endings of strong constraint sentences (Fig 2A). Thus, similarity was increased by higher levels of sentential constraint and of final word expectancy.

**Figure 2.**
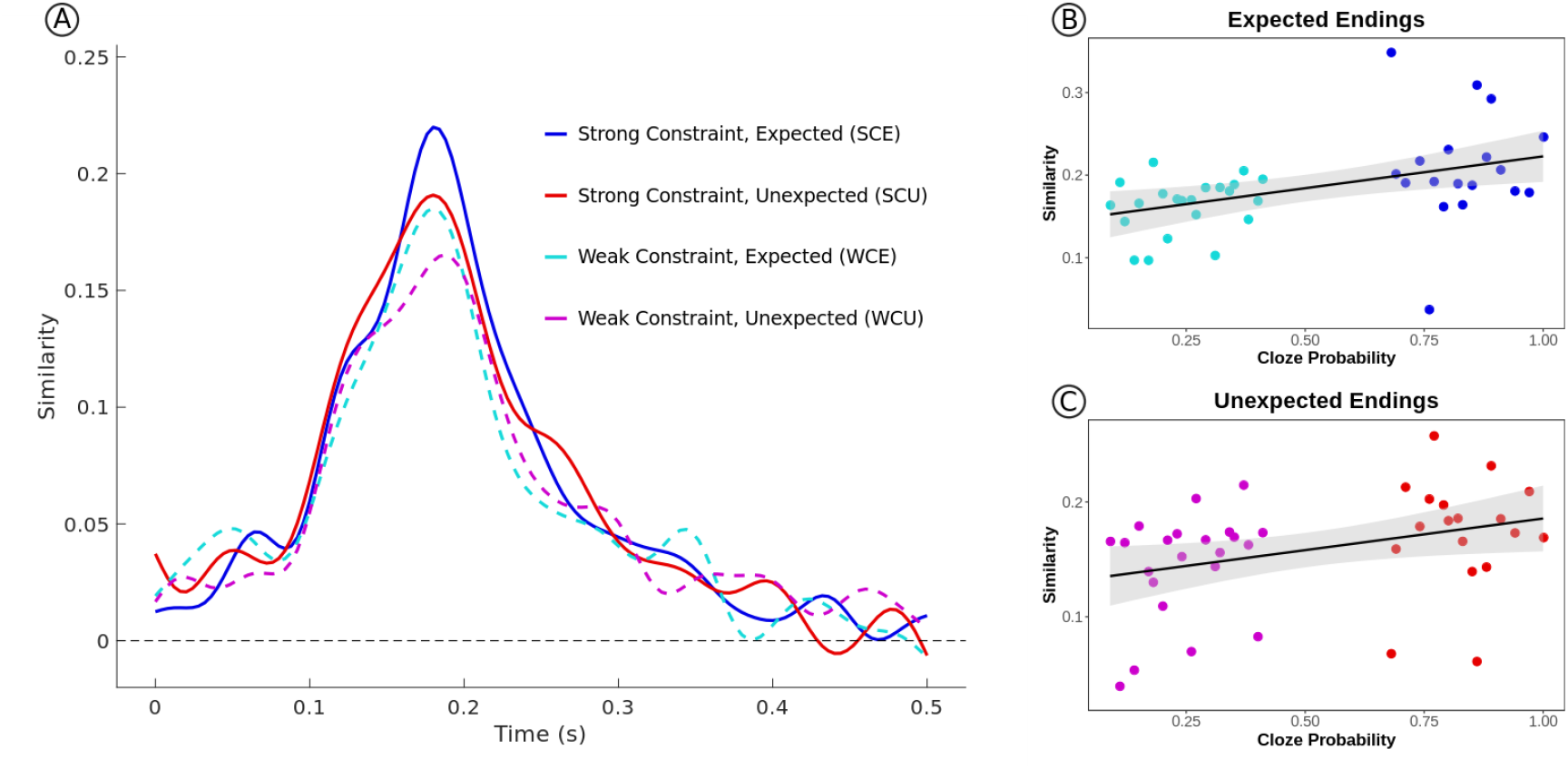
Results of the spatial RSA. **A)** The similarity time-course is shown for each of the four conditions. A peak in similarity is observed that varies with constraint and expectancy. **B)** The correlation between neural similarity and cloze probability for sentences with expected endings. **C)** The correlation between neural similarity and cloze probability of the expected sentence endings for sentences with unexpected endings. Both correlations are significantly positive.

Treating sentential constraint as a categorical variable is somewhat artificial, as cloze values range continuously. To more specifically test predictability’s effect on the observed neural similarity, an item-level analysis was performed. The similarity values were averaged across subjects for each item with the same cloze probability for the expected endings (combining SCE & WCE), and a linear regression was run predicting neural similarity from cloze probability. Cloze probability significantly predicted neural similarity for the expected endings (*t* = 2.91, *r*^*2*^ = 0.18, *p* < 0.01); see Fig 2B. Given that the response to unexpected endings was overall lower in similarity but showed sensitivity to constraint, an additional regression was run predicting neural similarity for the unexpected endings (combining SCU & WCU) from graded sentential constraint (i.e., the cloze probability of the most expected ending for that sentence). As seen in Fig 2C, there was a significant linear relationship (*t* = 2.24, *r*^*2*^ = 0.12, *p* = 0.03), suggesting that the prediction signal was graded with sentential constraint. Note that while the majority of the range of cloze probabilities was sampled with these stimuli, there were no items in the middle range of cloze probability (~40-60%). However, relationships between neural activity (e.g. N400s) and cloze probability are usually linear [31], and thus it is unlikely that the inclusion of middle-range cloze items would produce a nonlinear relationship.

The observed neural similarity pattern may have been driven by similarity in lexical characteristics of the pre-final and final words leading to overlap in patterns of activity on the scalp, not by pre-activation of the final word. To test this, we used linear mixed-effects models to predict pattern similarity with cloze probability and lexical characteristics on a single-trial level. For each trial, the average similarity derived from spatial RSA from 160-210 ms was extracted. The first analysis predicted trial-level similarity values from cloze probability for expected sentence endings, and a random intercept for participants was included. Given the hypothesis that lexical similarity could explain the observed effect, we created differences measures by taking the absolute value of the difference between the pre-final word frequency from the final word frequency, as well as the word length^1^, and included these measures in the model. Thus, the model was constructed as follows:

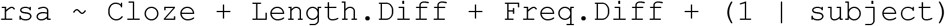

Random slopes for fixed effects of interests were not included due to issues with model convergence. Significance of fixed effects of the model was assessed with t-tests using the Satterthwaite method of approximation for degrees of freedom. The results of this analysis are reported in Table 1. Cloze probability remained a significant predictor of neural similarity, even with lexical characteristics included. Neither word length nor frequency were significant, though frequency trended toward significance.

**Table 1.** Fixed effect estimates and tests of significance for mixed effects model predicting trial level similarity values for expected endings derived from spatial RSA.

|  | Estimate | Std. Error | Dg. freedom | *t*-value | *p*-value |
| --- | --- | --- | --- | --- | --- |
| Intercept | 0.163 | 0.026 | 59.06 | 6.24 | <0.01* |
| Cloze | 0.048 | 0.016 | 3,932 | 3.04 | <0.01* |
| Length | 0.004 | 0.003 | 3,932 | 1.35 | 0.18 |
| Freq | -0.009 | 0.005 | 3,932 | -1.80 | 0.07 |

The second analysis focused on unexpected sentence endings. Here, the model was similar to the first model described previously; however, the cloze probability of the expected ending of the sentence was included instead of the cloze probability of the unexpected ending. Thus, the model was constructed as:

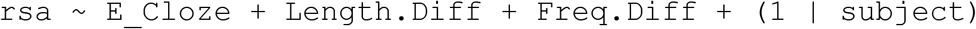

The fixed effects results are reported in Table 2. As before, cloze probability remained a significant predictor of similarity even after including lexical variables. However, word frequency was also a significant predictor of similarity. Thus, while differences in word frequency may have contributed to the effect for unexpected endings, the pre-activation account remains a viable explanation.

**Table 2.**
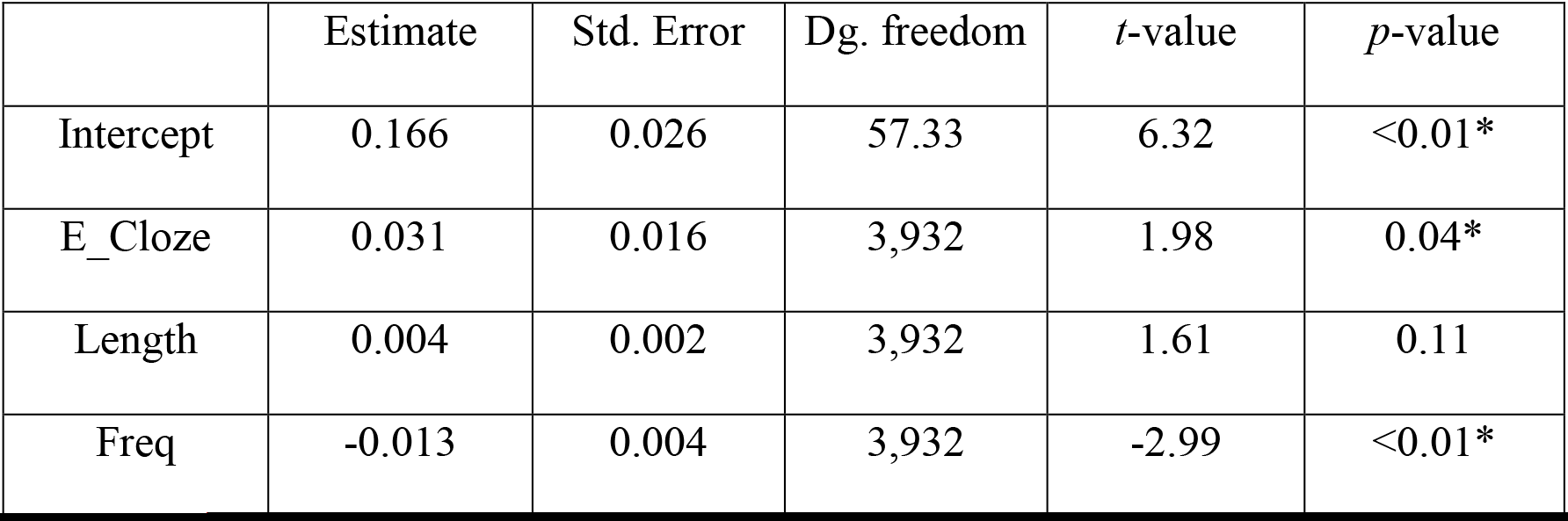
Fixed effect estimates and tests of significance for mixed effects model predicting trial level similarity values for unexpected endings derived from spatial RSA.

Predictions may have been generated or may be detectable prior to the onset of the pre-final word. To assess, we performed the same spatial RSA method comparing neural activity patterns of the final word to the word prior to the pre-final word (2 word positions back). A peak in pattern similarity was found in the same time window as the pre-final word analysis, but with lower magnitude (Figure 3). A repeated measures ANOVA run on similarity values as in the pre-final word analysis revealed no significant main effects (constraint, F_(1,30)_ = 0.14, *p* = 0.71; expectancy, F_(1,30)_ = 0.08, *p* = 0.78) or interaction (F_(1,30)_ = 3.69, *p* = 0.06). Additionally, a linear regression predicting pattern similarity from cloze probability was not significant for either expected (*t* = 1.65, *r*^*2*^ = 0.07, *p* = 0.11) or unexpected (*t* = 0.21, *r*^*2*^ < 0.01, *p* = 0.84) sentence endings. A spatial RSA comparing final word activity to words 3 word positions back also showed no significant effects of predictability (all p-values > 0.05; Figure 3). Thus, prediction-related pattern similarity differences were only reliable immediately prior to the onset of the sentence final word.

**Figure 3.**
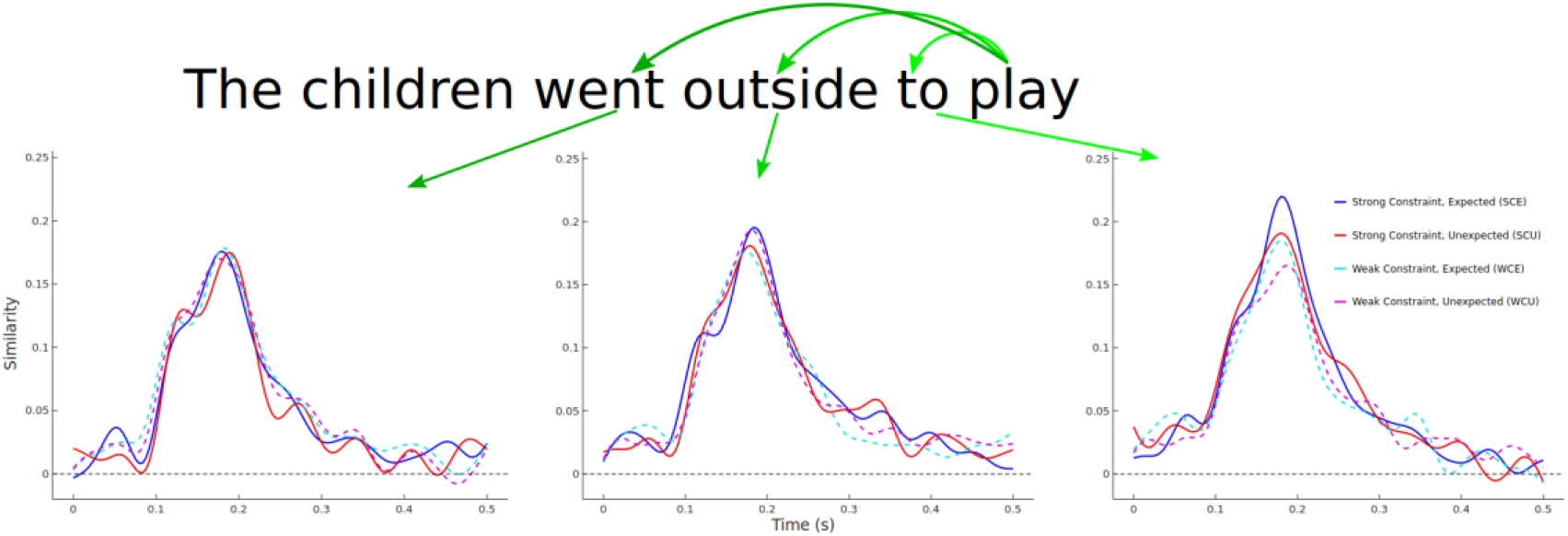
Spatial RSA for sentence final words and pre-final words at three different positions: immediately preceding (right plot), two words back (center plot), and three words back (left plot). Significant differences are observed only for pre-final words immediately preceding sentence final words.

### Generalization Analysis

To assess the fate of predicted representations across time, we employed a time generalization analysis. The pattern of activity across channels at each timepoint from 0 to 300 ms following the pre-final word was correlated with every timepoint from 0 to 500 ms following the final word. The resulting time × time matrices were analyzed by submitting pair-wise contrasts to cluster-based permutation tests.

Permutation tests revealed four clusters of interest that reached a significant cluster-wise threshold of *p* < 0.05 (Fig 4A). First, in the contrast of expected words (SCE - WCE), a positive cluster was found (pre-final word time: 150-275 ms; post-final word time: 160-265 ms; *p* < 0.01), with SCE similarity greater than WCE. This cluster likely reflects the same effect found with the initial spatial RSA analysis, and a similar positive cluster was found in the contrast of SCE and WCU words (pre-final word time: 150-250 ms; post-final word time: 155-210 ms; *p* = 0.01). These positive clusters demonstrate that early pre-final word and post-final word similarity is greater for more predictable sentence endings. Comparisons of other pairs did not yield significant positive clusters. It is important to note, however, that the cluster-based permutation tests are more statistically conservative than the analysis focused on the specific peak. Critically, even under a conservative analytical approach, similarity is found to be enhanced by both expectancy and constraint.

**Figure 4.**
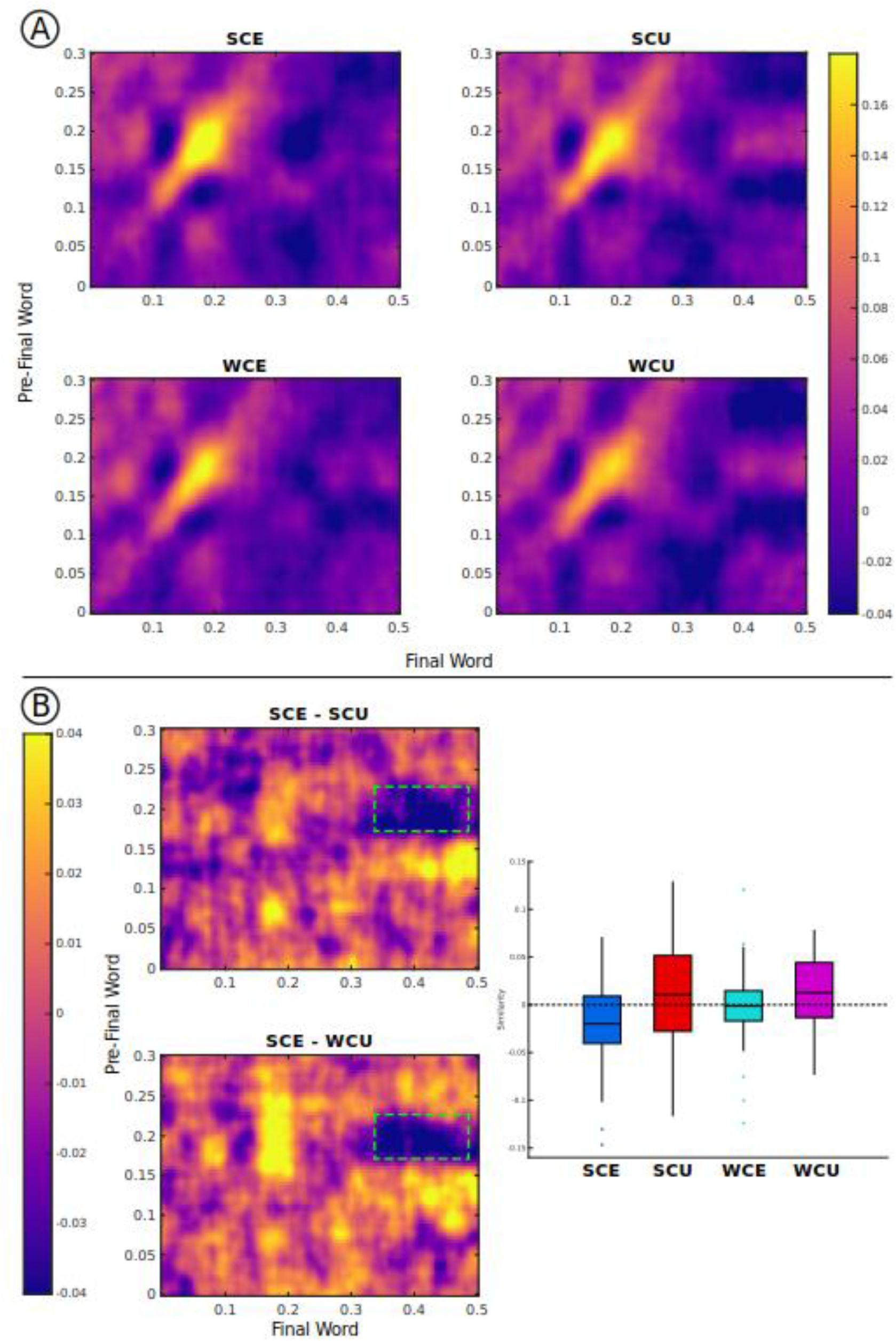
Results of the spatial generalization RSA. **A)** Generalization matrices for each condition are plotted. The color intensity represents neural similarity. The strong increase observed in each condition reflects the previously observed spatial RSA peak (Figure 2A). **B)** Differences in generalization matrices for SCE-SCU and SCE-WCU. The green box highlights the significant cluster found for both differences. The bar-plot displays similarity values extracted from this time window, with SCE similarity significantly below zero, demonstrating repulsion.

The permutation tests also identified two similar negative clusters, with SCE words showing reduced similarity compared to both SCU words (pre-final word time: 170-230 ms; post-final word time: 315-475 ms; *p* < 0.01) and WCU words (pre-final word time: 175-230 ms; post-final word time: 340-500 ms; *p* < 0.01). Note that this effect reflected similarity of pre-final word activity in the time window of the previously reported spatial RSA effect and post-final word activity in a later time window, roughly the span of the N400 component of the ERP. To examine this effect further, we performed exploratory post-hoc analyses. Follow-up analyses on extracted similarity values (pre-final word time: 175-230 ms; post-final word time: 340-475) showed that similarity for SCE words was reduced compared to all other conditions (WCE: *t*_(30)_ = −2.60, *p* = 0.01; SCU: *t*_(30)_ = −3.62, *p* < 0.01; WCU: *t*_(30)_ = −4.33, *p* < 0.01), and, in fact, was significantly less than 0 (*t*_(30)_ = −2.62, *p* = 0.01). This pattern is depicted in Figure 4B.

An alternative explanation of these results is that the similarity difference results reflect a confound of univariate activation magnitude, which has been shown to affect pattern similarity results in fMRI studies [60]. The observed clusters were in the time window of the N400 following the final word, which does show amplitude differences in a similar pattern to the reported similarity pattern. We performed an additional analysis to test for this possibility. RSA was used to compare activity elicited by sentence final words with activity elicited by pre-final sentence words from different sentences that were the same as the pre-final word of the same sentence. For instance, the sentence “Father carved the turkey with *a* knife” has the same pre-final word as “His touch was light as *a* feather”. Here, we measured the similarity of the activity from the word knife in the first sentence to activity from the word *a* in the second sentence. This was done for all pre-final words that matched the pre-final word in the sentence, and the resultant correlation time-courses across trials were averaged. This allowed us to compare similarity when the pre-final word and final word were exactly the same, but only the sentence differed. This means only the activity at the pre-word differed, as the univariate effect of the N400 at the final word was the same as in the original analysis,.

The results of this analysis are shown in Figure 5. The difference plots are the same as those in Figure 4B – however, the negative cluster is not apparent. Cluster-based permutation tests revealed no significant clusters for either condition. Thus, the results are unlikely to be driven by univariate confounds.

**Figure 5.**
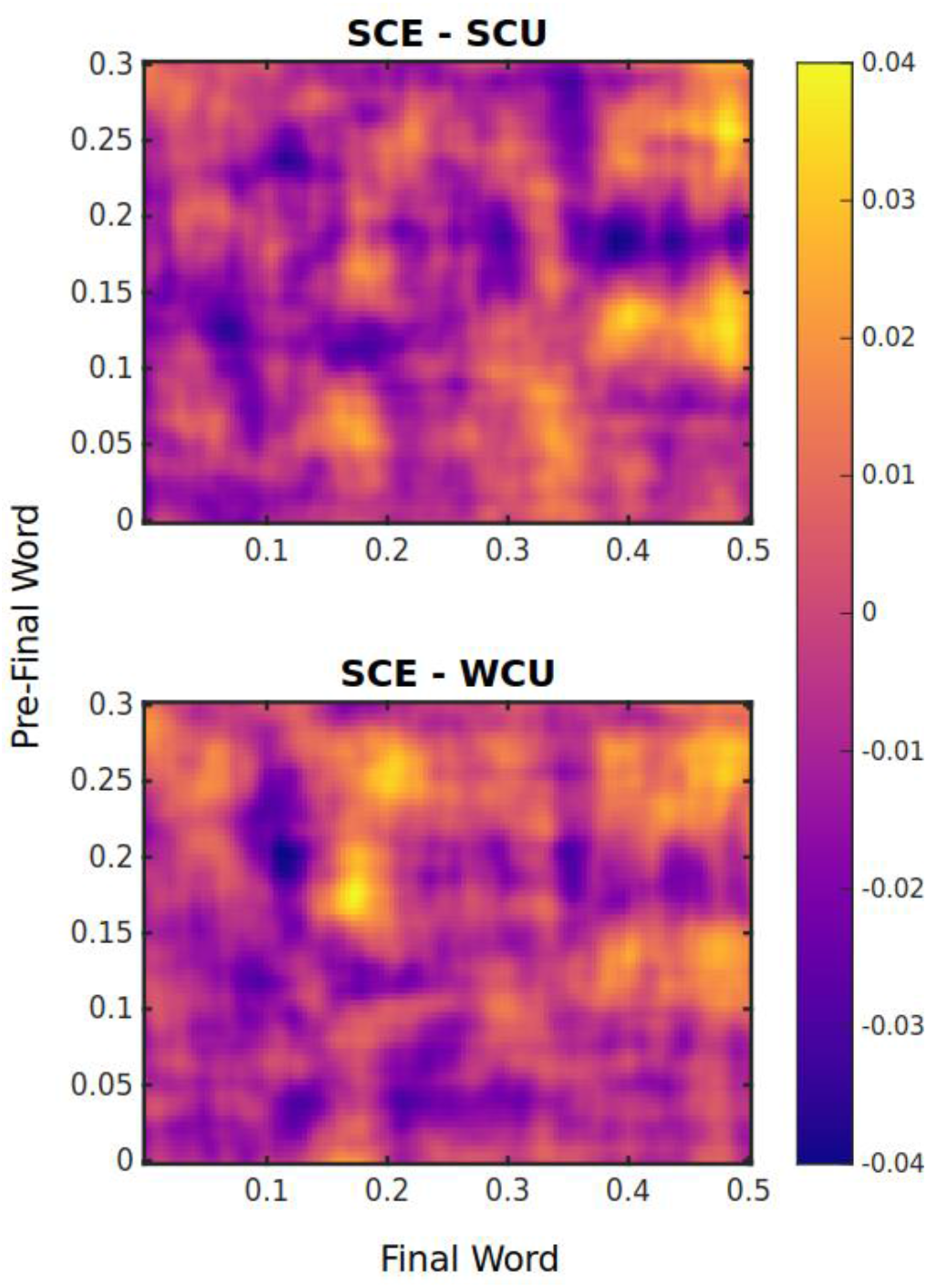
Results of the between-sentence spatial generalization analysis. The plots show differences between conditions (SCE-SCU for the top plot, SCE-WCU for the bottom plot). The later negative cluster found in the within-sentence generalization analysis is not observed.

### Effect topographies

To characterize the topography of the early spatial RSA effect, we used temporal RSA. Instead of correlating signals at each time-point, this analysis correlated the time series of pre-final word and final word activity in the selected window of analysis at each sensor on the scalp. Note that spatial RSA relates spatial patterns across time, whereas temporal RSA relates temporal patterns across space, and thus the statistical pattern of these results may differ. Temporal RSA showed that similarity was greatest over occipital channels across all conditions (Fig 6).

**Figure 6.**
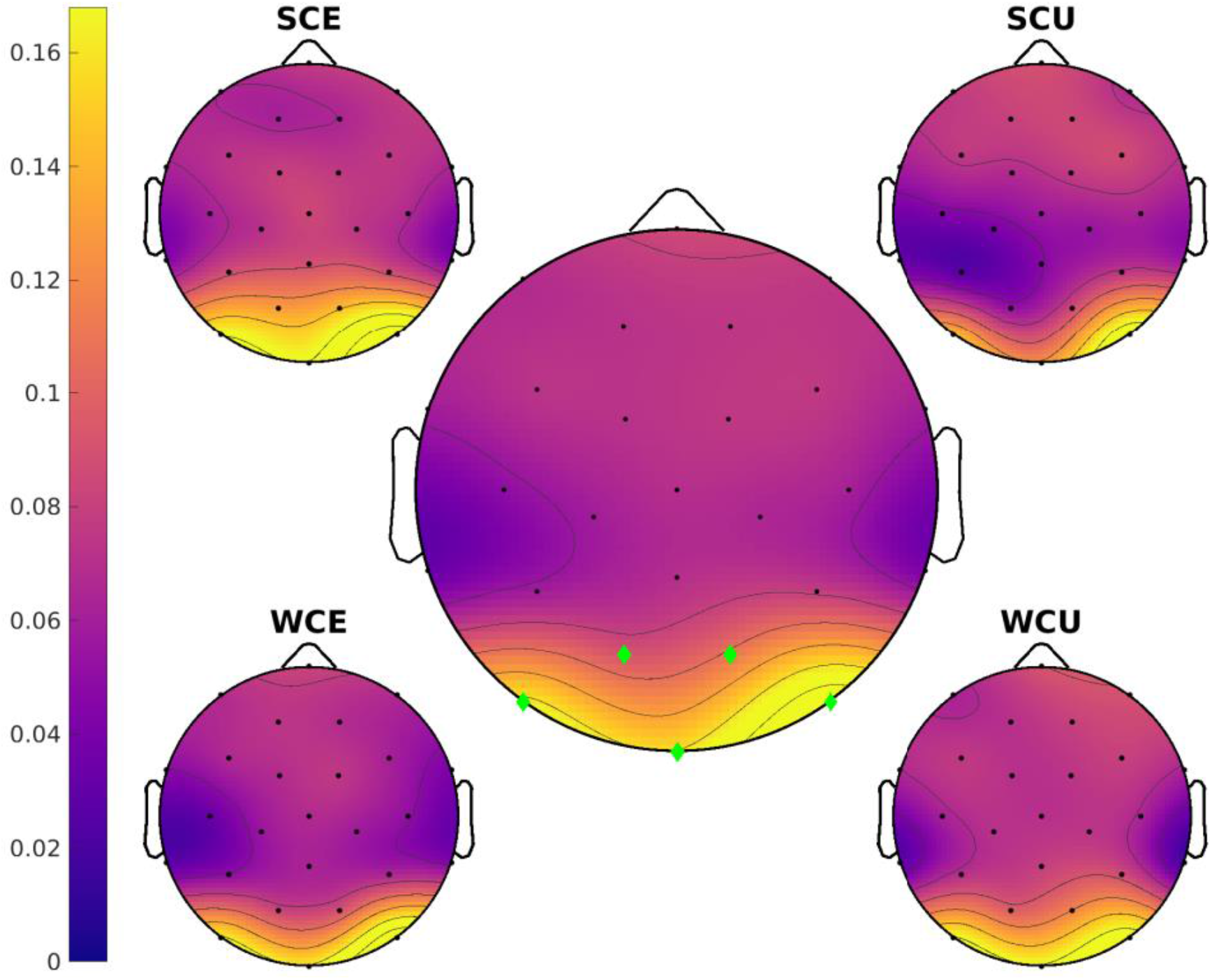
Temporal RSA results for the spatial RSA peak. The central topography plot shows the average across conditions, with the channels with the largest similarity values as green diamonds. The topography for each condition is also plotted. A strong occipital topography is observed.

Note that the topographies for each of the four conditions did not significantly differ from one another. However, this analysis is focused on the similarity in time at each channel, not the similarity across channels at each time-point. Thus, the resultant topography plots highlight the channels where the pre-final and final word time-series were the most similar. It is thus not surprising that the four conditions would not differ in this analysis, as they are likely to all reflect the same process, which varies in degree with prediction strength and level of match between the prediction and the input.

We performed a similar analysis to characterize the late negative cluster found in the time generalization analysis. We implemented a sliding window approach, in which the time series of pre-final word activity from 170-230 ms was correlated with final word activity from 340-480 ms across successive 60 ms windows (Fig 7). Note that these time windows were designated by identifying the minimum and maximum time values of the significant clusters reported previously. An initial dissimilarity was observed over occipital channels, which was more pronounced and sustained for SCE words. This was followed by an increase in similarity over central channels for unexpected words, but not for expected words. This timecourse corroborates the results from the cluster analyses; namely, strongly expected words show less similarity later in time compared to less expected words.

**Figure 7.**
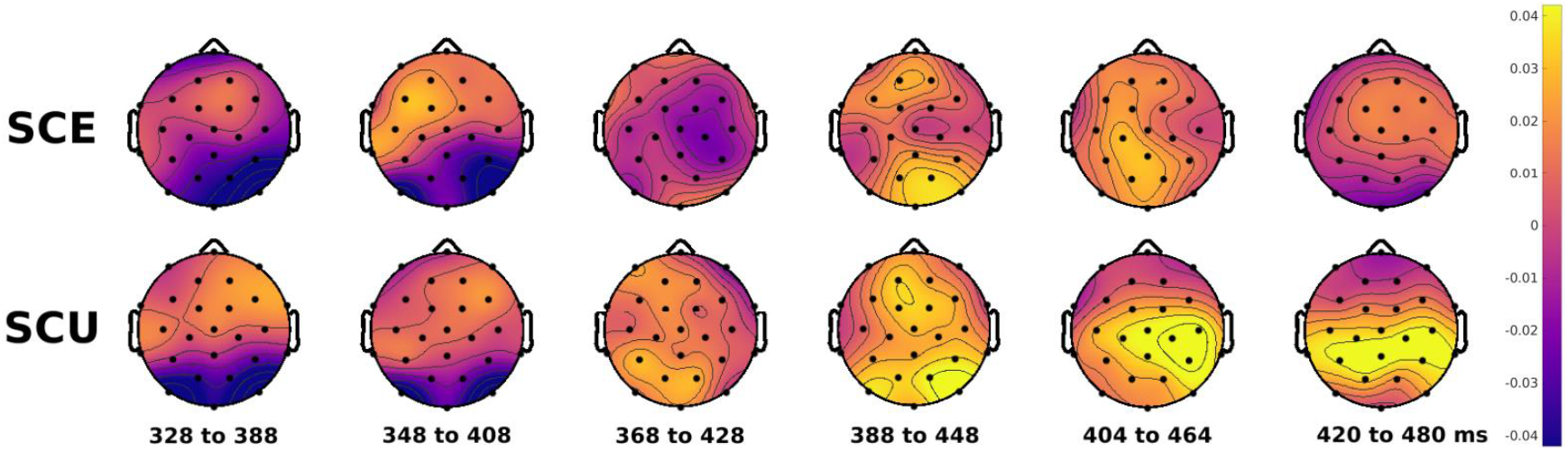
Sliding window temporal RSA results for the late cluster. Similarity topographies are shown at different time windows, where the activity from the final word in the displayed time window is correlated with pre-final word activity from 170-230 ms. SCE shows broad dissimilarity that stays near zero over time. SCU shows early dissimilarity, followed by positive similarity over posterior channels.

## Discussion

Numerous studies have investigated the neural consequences of predictability during language comprehension, but the specific mechanisms, timing, and extent of anticipatory pre-activation have remained elusive. Here, we used RSA to compare patterns of neural activity prior to a sentence-final word to the activity following a sentence-final word, allowing us to examine the timing of generating predictions, as well as their specificity. Our results provide persuasive evidence that people predict upcoming information probabilistically and that those predictions affect word processing quite rapidly. Additionally, predictions appear to be generated or allocated at specific times, i.e. close in time to the upcoming final word. Finally, we provide novel evidence that activity patterns representing pre-activations are “repulsed” following confirmation of predictions. Together, these results elucidate the neural mechanisms of prediction during the comprehension of language and potentially provide insight into general mechanisms of prediction in the brain.

Spatial RSA revealed an increase in neural similarity between pre-final word activity and final word activity that extended from approximately 100-300 ms following pre-final word onset. This similarity was graded with the cloze probability of the sentence final word, such that similarity decreased as the word became less predictable, showing that predictions are graded. This relationship between neural similarity and cloze probability may reflect a graded degree of effort, in which neural resources are allocated towards anticipatory processing and the level of resource allocation is dependent on predictability and, thus, a function of constraint. Alternatively, this pattern may reflect a graded degree of success, in which the probability of a match between the predicted and actual outcome is greater with higher levels of sentential constraint. Additionally, we found that similarity was greatly reduced for unexpected but semantically plausible final words compared to expected final words, suggesting that the pre-activated features were at some level *specific* to the expected word. Thus, we provide the first results that, when reading language, features of specific upcoming words are rapidly pre-activated prior to their onset, and the magnitude of this anticipatory processing is graded with predictability.

Examining pattern similarity of the sentence final word and words prior to the pre-final word revealed that a significant relationship between similarity and predictability was present only for the pre-final word. In other words, evidence of anticipatory pre-activation was found only immediately prior to the word being predicted. This finding could reflect that predictions were generated rapidly following the onset of the pre-final word, and not before. Alternatively, the observed signal may not reflect the time at which information became available to the system, but, instead, the time at which the production system allocated resources towards explicitly forming a prediction of a particular type [61]. In a recent study, neural pre-activation of a series of expected simple visual stimuli occurred in visual cortex only after the first stimulus in the train was presented [62]; thus, there is precedent that the pre-final word could serve as a cue for pre-activation of visual features of the upcoming final word. In the current study, a sentence like “The bad boy was sent to his room” may have allowed some level of anticipation of the final word “room” even at the time of the word “sent” based on the semantics of the sentence. However, as previously described, the contents of prediction are multifaceted in nature, and these different features may be generated at different times. Sentence level semantic predictions may manifest in different neural signals than those observed here, and may influence the predictions generated at other levels, e.g. orthographic or syntactic. We do not claim that the pre-activation signal reported here reflects pre-activation of all linguistic information; there are likely other signals left to be identified relating to other levels of prediction. What our data reveal is that some aspects of prediction – possibly, as discussed next, prediction of specific word forms – seem to be specifically timed, perhaps cued by, for example, the preceding (usually function) word suggesting the imminent arrival of the target noun. This process could be epiphenomenal, i.e. stimuli automatically lead to pre-activation of features of associated stimuli that may follow. Alternatively, such a mechanism could be tailored by the nervous system to be beneficial for efficient behavior during language processing; for example, cued pre-activation could guide eye movement behavior during reading to reduce reading times and/or skip over easily predicted information [22].

Temporal RSA revealed an occipital topography in the time window of the pre-activation. Given EEG’s limited spatial resolution, it is difficult to pinpoint the exact networks that were involved in the observed pre-activation. However, recent related work may provide insight into the neural systems involved in anticipatory processing. For instance, a recent intracranial EEG study with temporal lobe epileptic patients used a similar method as used here to compare high-frequency hippocampal activity before and after naming predictable or unpredictable pictures [63]. Hippocampal activity was more similar when the picture was more predictable, mirroring the findings of this study. Given that the hippocampus plays a central role in several aspects of language processing [64], statistical learning [65], as well as predicting and imagining the future [66–67], it seems reasonable to think it could play a role in predicting upcoming linguistic features.

One proposed mechanism is that the hippocampus coordinates sensory pre-activation of upcoming information through a pattern completion process [68–69]. This proposal is in line with MEG results from a picture-word matching paradigm that demonstrated prediction of visual word form features [41]. Here, MEG source localization of activity prior to predicted target words revealed left medial temporal and occipital sources, with temporal activity slightly preceding occipital activity. Thus, temporal structures could pre-activate lexical information, leading to pre-activation of form features in sensory cortex. Our combined results – the timing of the similarity effect, as well as the strongly occipital topography – are consistent with this account, and suggest the increased similarity may have reflected overlap of pre-activated and observed lower level orthographic lexical features.

Consistent with the idea that the observed RSA signal might reflect prediction of word orthography, we observed that, although the pre-final and final word similarity was reduced for unexpected sentence completions, it was not abolished, and it varied with constraint. This differs from the pattern of semantic-based facilitation seen on the N400, wherein unexpected sentence completions elicit large N400s that do not differ by constraint [24]. Although, by design, there was likely little to no semantic overlap between expected and unexpected completions, orthographic space is more constrained, such that even unexpected words are likely to sometimes carry expected low-level features (shared length, a shared letter, etc.). The observed RSA pattern suggests that, although the unexpected endings were not predicted (and thus globally less similar), there was, in some cases, some level of – likely orthographic -- featural overlap. This hypothesis could be tested further by examining neural similarity to unexpected items that are specifically designed to be orthographically similar but semantically dissimilar to expected words.

The constraint-sensitivity of the similarity response for unexpected words shows that the prediction signal itself was variable in strength and/or fidelity, based on the predictability of the upcoming final word, an idea consistent with a probabilistic prediction account [1,5,70]. More weakly constraining sentences, by their nature, permit a wider range of completions, both at the semantic and orthographic level. Thus, if prediction were ubiquitous, the possibility of a match, at any level of analysis, would tend to be higher under weak constraint; yet similarity was reduced for unexpected items in weakly compared to more strongly constraining contexts. Rather than predicting to the same degree and/or level every time, the brain may essentially utilize a generative model to rationally and optimally allocate resources towards anticipatory processing based on the available contextual evidence and projected costs of pre-activation. A similar mechanism has been proposed to explain generalization and adaptation during speech perception [71]. If we recognize a familiar speaker, we may generate more specific predictions or activate more features of upcoming utterances from this speaker compared to an unfamiliar one. Here, individuals may have generated more specific predictions – perhaps including orthographic features – when the sentential context was highly biased toward a particular outcome. Since the pre-activation occurred prior to the final word, the representational similarity was graded by cloze probability even for unexpected sentence endings, as the representation of the final word was weaker when predictability was lower, leading to less of a possibility of even incidental featural overlap. To our knowledge, this result is one of the first to demonstrate rapid probabilistic pre-activation. Additional work using this method may shed light on the debate between serial and parallel predictions; namely, whether similarity for other possible sentence endings varies with their completion probability, or a graded response is only found for a single word representation.

To further probe the fate of predicted representations, we conducted a time generalization analysis, in which the spatial pattern of neural activity at each time point following the pre-final word was correlated with the pattern at each time point following the final word. This revealed a surprising finding: Early pre-final word activity was less similar to later final word activity for strongly constrained expected endings compared to unexpected endings, and in fact had negative similarity or anti-correlation. Anti-correlation has only rarely been observed in other studies utilizing RSA but has been found in episodic memory studies; namely, hippocampal firing patterns representing events occurring in different contexts are anti-correlated [72]. Similarly, hippocampal representations of overlapping spatial routes become anti-correlated, or demonstrate “repulsion” or “differentiation”, over time [73]. Here, final word activity became differentiated from pre-final word activity after the early increase in similarity. The fact that this occurred most in cases wherein strong predictions were formed and confirmed suggests that the representation of the pre-final word was not repulsed, but in fact, the features of the final word that were pre-activated were repulsed. Such a result is in line with other findings that, downstream, individuals have impoverished representations and impaired memory for predicted words [49–50]. This provides a view of anticipatory processing during language comprehension in which prediction allows for rapid verification of incoming words, leading to more efficient processing in the moment, but, once the predicted information is verified, the brain shifts processing away, essentially “leaving behind” the predicted information.

Another insight from the time generalization analysis is that the similarity increase during the pre-final period was not sustained over the delay prior to the onset of the word, as might have been predicted by analogy to some accounts of the maintenance of information in working memory [74]. Indeed, maintained neural firing to sustain predictions would seem a highly inefficient and metabolically costly strategy for the brain, and thus not likely to be the mechanism of pre-activation. Even sustained working memory signals are observed after averaging many trials; in actuality, spiking during delay periods on single trials is sparse and varies in time [75]. More recent evidence suggests that activity at specific frequencies coordinates in bursts to produce rapid synaptic weight changes, which efficiently code information [76–77]. A similar mechanism seems likely to be utilized for rapid predictive coding and would explain how a lack of maintained delay-related activity could still produce pre-activation of upcoming information.

As a set, our findings demonstrate that the brain rapidly pre-activates specific features of upcoming words during language comprehension and does so in a timely manner to allow for efficient processing. These results not only conclusively demonstrate that individuals utilize prediction during comprehension, but also shed light on one of the fundamental functions of the brain. Environmental cues may lead to the generation of predictions of associated information or stimuli, and, once the pre-activated stimulus is encountered and confirmed, the brain may rapidly shift processing away to focus resources on other objectives.

## Author Contributions

R.J.H. designed and performed the analyses. R.J.H. and K.D.F. wrote the paper together.

## Competing Interests

The authors declare no competing financial interests.

## Data Availability

For protection of participants’ privacy, the data is available upon request to the authors. Please email the corresponding author for more information.

## Footnotes

1 Orthographic neighborhood size is also a lexical variable known to affect electrophysiological responses to words. However, word length is highly correlated with orthographic neighborhood. To simplify the model, we only included word length and frequency.

## Notes

### Competing Interest Statement

The authors have declared no competing interest.

## References

1. Friston, K., & Kiebel, S. (2009). Predictive coding under the free-energy principle. Philosophical Transactions of the Royal Society B: Biological Sciences, 364(1521), 1211–1221. https://doi.org/10.1098/rstb.2008.0300

2. Huang, Y., & Rao, R. P. N. (2011). Predictive coding. Wiley Interdisciplinary Reviews: Cognitive Science, 2(5), 580–593. https://doi.org/10.1002/wcs.142

3. Kiebel, S. J., Daunizeau, J., & Friston, K. J. (2009). Perception and hierarchical dynamics. Frontiers in Neuroinformatics, 3. https://doi.org/10.3389/neuro.11.020.2009

4. Federmeier, K. D. (2007). Thinking ahead: The role and roots of prediction in language comprehension. Psychophysiology, 44(4), 491–505. https://doi.org/10.1111/j.1469-8986.2007.00531.x

5. Kuperberg, G. R., & Jaeger, T. F. (2016). What do we mean by prediction in language comprehension? Language, Cognition and Neuroscience, 31(1), 32–59. https://doi.org/10.1080/23273798.2015.1102299

6. Kutas, M., DeLong, K. A., & Smith, N. J. (2011). A look around at what lies ahead: Prediction and predictability in language processing. In M. Bar, Predictions in the brain: Using our past to generate a future (pp. 190–207). Oxford University Press.

7. Pickering, M. J., & Gambi, C. (2018). Predicting while comprehending language: A theory and review. Psychological Bulletin, 144(10), 1002–1044. https://doi.org/10.1037/bul0000158

8. Kim, A., & Lai, V. (2012). Rapid interactions between lexical semantic and word form analysis during word recognition in context: Evidence from ERPs. Journal of Cognitive Neuroscience, 24(5), 1104–1112. https://doi.org/10.1162/jocn_a_00148

9. Laszlo, S., & Federmeier, K. D. (2009). A beautiful day in the neighborhood: An event-related potential study of lexical relationships and prediction in context. Journal of Memory and Language, 61(3), 326–338. https://doi.org/10.1016/j.jml.2009.06.004

10. DeLong, K. A., Urbach, T. P., & Kutas, M. (2005). Probabilistic word pre-activation during language comprehension inferred from electrical brain activity. Nature Neuroscience, 8(8), 1117–1121. https://doi.org/10.1038/nn1504

11. Vissers, C. Th. W. M., Chwilla, D. J., & Kolk, H. H. J. (2006). Monitoring in language perception: The effect of misspellings of words in highly constrained sentences. Brain Research, 1106(1), 150–163. https://doi.org/10.1016/j.brainres.2006.05.012

12. Federmeier, K. D., & Kutas, M. (1999). A rose by any other name: Long-term memory structure and sentence processing. Journal of Memory and Language, 41(4), 469–495. https://doi.org/10.1006/jmla.1999.2660

13. Lau, E. F., Holcomb, P. J., & Kuperberg, G. R. (2013). Dissociating N400 effects of prediction from association in single-word contexts. Journal of Cognitive Neuroscience, 25(3), 484–502. https://doi.org/10.1162/jocn_a_00328

14. Dikker, S., Rabagliati, H., Farmer, T. A., & Pylkkänen, L. (2010). Early occipital sensitivity to syntactic category is based on form typicality. Psychological Science, 21(5), 629–634. https://doi.org/10.1177/0956797610367751

15. Van Berkum, J. J. A., Brown, C. M., Zwitserlood, P., Kooijman, V., & Hagoort, P. (2005). Anticipating upcoming words in discourse: Evidence from erps and reading times. Journal of Experimental Psychology: Learning, Memory, and Cognition, 31(3), 443–467. https://doi.org/10.1037/0278-7393.31.3.443

16. Huettig, F., & Guerra, E. (2019). Effects of speech rate, preview time of visual context, and participant instructions reveal strong limits on prediction in language processing. Brain Research, 1706, 196–208. https://doi.org/10.1016/j.brainres.2018.11.013

17. Wlotko, E. W., & Federmeier, K. D. (2015). Time for prediction? The effect of presentation rate on predictive sentence comprehension during word-by-word reading. Cortex, 68, 20–32. https://doi.org/10.1016/j.cortex.2015.03.014

18. Hess, D. J., Foss, D. J., & Carroll, P. (1995). Effects of global and local context on lexical processing during language comprehension. Journal of Experimental Psychology: General, 124(1), 62–82. https://doi.org/10.1037/0096-3445.124.1.62

19. Kleiman, G. M. (1980). Sentence frame contexts and lexical decisions: Sentence-acceptability and word-relatedness effects. Memory & Cognition, 8(4), 336–344. https://doi.org/10.3758/BF03198273

20. Schwanenflugel, P. J., & LaCount, K. L. (1988). Semantic relatedness and the scope of facilitation for upcoming words in sentences. Journal of Experimental Psychology: Learning, Memory, and Cognition, 14(2), 344–354. https://doi.org/10.1037/0278-7393.14.2.344

21. Altmann, G. T. M., & Kamide, Y. (1999). Incremental interpretation at verbs: Restricting the domain of subsequent reference. Cognition, 73(3), 247–264. https://doi.org/10.1016/S0010-0277(99)00059-1

22. Ehrlich, S. F., & Rayner, K. (1981). Contextual effects on word perception and eye movements during reading. Journal of Verbal Learning and Verbal Behavior, 20(6), 641–655. https://doi.org/10.1016/S0022-5371(81)90220-6

23. Staub, A., & Clifton, C. (2006). Syntactic prediction in language comprehension: Evidence from either…or. Journal of Experimental Psychology: Learning, Memory, and Cognition, 32(2), 425–436. https://doi.org/10.1037/0278-7393.32.2.425

24. Federmeier, K. D., Wlotko, E. W., De Ochoa-Dewald, E., & Kutas, M. (2007). Multiple effects of sentential constraint on word processing. Brain Research, 1146, 75–84. https://doi.org/10.1016/j.brainres.2006.06.101

25, Szewczyk, J. M., & Schriefers, H. (2018). The N400 as an index of lexical preactivation and its implications for prediction in language comprehension. Language, Cognition and Neuroscience, 33(6), 665–686. https://doi.org/10.1080/23273798.2017.1401101

26. Thornhill, D. E., & Van Petten, C. (2012). Lexical versus conceptual anticipation during sentence processing: Frontal positivity and N400 ERP components. International Journal of Psychophysiology, 83(3), 382–392. https://doi.org/10.1016/j.ijpsycho.2011.12.007

27. Dikker, S., & Pylkkanen, L. (2011). Before the N400: Effects of lexical–semantic violations in visual cortex. Brain and Language, 118(1-2), 23–28. https://doi.org/10.1016/j.bandl.2011.02.006

28. Wang, L., Hagoort, P., & Jensen, O. (2018). Language prediction is reflected by coupling between frontal gamma and posterior alpha oscillations. Journal of Cognitive Neuroscience, 30(3), 432–447. https://doi.org/10.1162/jocn_a_01190

29. Wang, L., Kuperberg, G., & Jensen, O. (2018). Specific lexico-semantic predictions are associated with unique spatial and temporal patterns of neural activity. ELife, 7, e39061. https://doi.org/10.7554/eLife.39061

30. DeLong, K. A., Quante, L., & Kutas, M. (2014). Predictability, plausibility, and two late ERP positivities during written sentence comprehension. Neuropsychologia, 61, 150–162. https://doi.org/10.1016/j.neuropsychologia.2014.06.016

31. Wlotko, E. W., & Federmeier, K. D. (2012). So that’s what you meant! Event-related potentials reveal multiple aspects of context use during construction of message-level meaning. NeuroImage, 62(1), 356–366. https://doi.org/10.1016/j.neuroimage.2012.04.054

32. Szewczyk, J. M., & Schriefers, H. (2013). Prediction in language comprehension beyond specific words: An ERP study on sentence comprehension in Polish. Journal of Memory and Language, 68(4), 297–314. https://doi.org/10.1016/j.jml.2012.12.002

33. Wicha, N. Y. Y., Moreno, E. M., & Kutas, M. (2003). Expecting gender: An event related brain potential study on the role of grammatical gender in comprehending a line drawing within a written sentence in spanish. Cortex, 39(3), 483–508. https://doi.org/10.1016/S0010-9452(08)70260-0

34. Nieuwland, M. S. et al., (2018). Large-scale replication study reveals a limit on probabilistic prediction in language comprehension. ELife, 7, e33468. https://doi.org/10.7554/eLife.33468

35. Nicenboim, B., Vasishth, S., & Rösler, F. (2019). Are words pre-activated probabilistically during sentence comprehension? Evidence from new data and a Bayesian random-effects meta-analysis using publicly available data. Preprint at https://psyarxiv.com/2atrh/

36. Roll, M., Söderström, P., Frid, J., Mannfolk, P., & Horne, M. (2017). Forehearing words: Pre-activation of word endings at word onset. Neuroscience Letters, 658, 57–61. https://doi.org/10.1016/j.neulet.2017.08.030

37. Söderström, P., Horne, M., Frid, J., & Roll, M. (2016). Pre-activation negativity (PrAN) in brain potentials to unfolding words. Frontiers in Human Neuroscience, 10. https://doi.org/10.3389/fnhum.2016.00512

38. Freunberger, D., & Roehm, D. (2017). The costs of being certain: Brain potential evidence for linguistic preactivation in sentence processing: Brain-potential evidence for linguistic preactivation. Psychophysiology, 54(6), 824–832. https://doi.org/10.1111/psyp.12848

39. Maess, B., Mamashli, F., Obleser, J., Helle, L., & Friederici, A. D. (2016). Prediction signatures in the brain: Semantic pre-activation during language comprehension. Frontiers in Human Neuroscience, 10. https://doi.org/10.3389/fnhum.2016.00591

40. León-Cabrera, P., Rodríguez-Fornells, A., & Morís, J. (2017). Electrophysiological correlates of semantic anticipation during speech comprehension. Neuropsychologia, 99, 326–334. https://doi.org/10.1016/j.neuropsychologia.2017.02.026

41. Dikker, S., & Pylkkänen, L. (2013). Predicting language: MEG evidence for lexical preactivation. Brain and Language, 127(1), 55–64. https://doi.org/10.1016/j.bandl.2012.08.004

42. Cichy, R. M., & Pantazis, D. (2017). Multivariate pattern analysis of MEG and EEG: A comparison of representational structure in time and space. NeuroImage, 158, 441–454. https://doi.org/10.1016/j.neuroimage.2017.07.023

43. Kriegeskorte, N., & Kievit, R. A. (2013). Representational geometry: Integrating cognition, computation, and the brain. Trends in Cognitive Sciences, 17(8), 401–412. https://doi.org/10.1016/j.tics.2013.06.007

44. Kriegeskorte, N., Mur, M., & Bandettini, P. A. (2008). Representational similarity analysis—Connecting the branches of systems neuroscience. Frontiers in Systems Neuroscience, 2. https://doi.org/10.3389/neuro.06.004.2008

45. Cichy, R. M., Ramirez, F. M., & Pantazis, D. (2015). Can visual information encoded in cortical columns be decoded from magnetoencephalography data in humans? NeuroImage, 121, 193–204. https://doi.org/10.1016/j.neuroimage.2015.07.011

46. Rommers, J., & Federmeier, K. D. (2018a). Lingering expectations: A pseudo-repetition effect for words previously expected but not presented. NeuroImage, 183, 263–272. https://doi.org/10.1016/j.neuroimage.2018.08.023

47. Heikel, E., Sassenhagen, J., & Fiebach, C. J. (2018). Time-generalized multivariate analysis of EEG responses reveals a cascading architecture of semantic mismatch processing. Brain and Language, 184, 43–53. https://doi.org/10.1016/j.bandl.2018.06.007

48. King, J-R., & Dehaene, S. (2014). Characterizing the dynamics of mental representations: The temporal generalization method. Trends in Cognitive Sciences, 18(4), 203–210. https://doi.org/10.1016/j.tics.2014.01.002

49. Rommers, J., & Federmeier, K. D. (2018b). Predictability’s aftermath: Downstream consequences of word predictability as revealed by repetition effects. Cortex, 101, 16–30. https://doi.org/10.1016/j.cortex.2017.12.018

50. Hubbard, R. J., Rommers, J., Jacobs, C. L., & Federmeier, K. D. (2019). Downstream behavioral and electrophysiological consequences of word prediction on recognition memory. Frontiers in Human Neuroscience, 13. https://doi.org/10.3389/fnhum.2019.00291

51. Bates, D., Mächler, M., Bolker, B., & Walker, S. (2015). Fitting Linear Mixed-Effects Models Using lme4. Journal of Statistical Software, 67(1), 1–48. https://doi.org/10.18637/jss.v067.i01

52. Kuznetsova, A., Brockhoff, P. B., & Christensen, R. H. (2017). lmerTest Package: Tests in Linear Mixed Effects Models. Journal of Statistical Software, 82(1), 1–26. https://doi.org/10.18637/jss.v082.i13

53. Tanner, D., Morgan-Short, K., & Luck, S. J. (2015). How inappropriate high‐ pass filters can produce artifactual effects and incorrect conclusions in ERP studies of language and cognition. Psychophysiology, 52(8), 997–1009. https://doi.org/10.1111/psyp.12437

54. Palmer, J. A., Kreutz-Delgado, K., & Makeig, S. (2012). AMICA: An adaptive mixture of independent component analyzers with shared components. Swartz Center for Computatonal Neursoscience, University of California San Diego, Tech. Rep.

55. Dimsdale-Zucker, H. R., & Ranganath, C. (2018). Representational similarity analyses: A practical guide for functional MRI applications. In Handbook of Behavioral Neuroscience (Vol. 28, pp. 509–525). Elsevier.

56. Ciuparu, A., & Mureşan, R. C. (2016). Sources of bias in single-trial normalization procedures. European Journal of Neuroscience, 43(7), 861–869. https://doi.org/10.1111/ejn.13179

57. Grandchamp, R., & Delorme, A. (2011). Single-trial normalization for event-related spectral decomposition reduces sensitivity to noisy trials. Frontiers in Psychology, 2, 236–236.

58. Luck, S. J., & Gaspelin, N. (2017). How to get statistically significant effects in any ERP experiment (And why you shouldn’t). Psychophysiology, 54(1), 146–157. https://doi.org/10.1111/psyp.12639

59. Maris, E., & Oostenveld, R. (2007). Nonparametric statistical testing of EEG- and MEG-data. Journal of Neuroscience Methods, 164(1), 177–190. https://doi.org/10.1016/j.jneumeth.2007.03.024

60. Coutanche, M. N. (2013). Distinguishing multi-voxel patterns and mean activation: Why, how, and what does it tell us? Cognitive, Affective, & Behavioral Neuroscience, 13(3), 667–673. https://doi.org/10.3758/s13415-013-0186-2

61. Dell, G. S., & Chang, F. (2014). The P-chain: Relating sentence production and its disorders to comprehension and acquisition. Philosophical Transactions of the Royal Society B: Biological Sciences, 369(1634), 20120394. https://doi.org/10.1098/rstb.2012.0394

62. Ekman, M., Kok, P., & de Lange, F. P. (2017). Time-compressed preplay of anticipated events in human primary visual cortex. Nature Communications, 8(1), 1–9. https://doi.org/10.1038/ncomms15276

63. Jafarpour, A., Piai, V., Lin, J. J., & Knight, R. T. (2017). Human hippocampal pre-activation predicts behavior. Scientific Reports, 7(1), 1–9. https://doi.org/10.1038/s41598-017-06477-5

64. Duff, M. C., & Brown-Schmidt, S. (2012). The hippocampus and the flexible use and processing of language. Frontiers in Human Neuroscience, 6. https://doi.org/10.3389/fnhum.2012.00069

65. Schapiro, A. C., Turk‐Browne, N. B., Norman, K. A., & Botvinick, M. M. (2016). Statistical learning of temporal community structure in the hippocampus. Hippocampus, 26(1), 3–8. https://doi.org/10.1002/hipo.22523

66. Buckner, R. L. (2010). The role of the hippocampus in prediction and imagination. Annual Review of Psychology, 61, 27–48. https://doi.org/10.1146/annurev.psych.60.110707.163508

67. Hassabis, D., Kumaran, D., Vann, S. D., & Maguire, E. A. (2007). Patients with hippocampal amnesia cannot imagine new experiences. Proceedings of the National Academy of Sciences, 104(5), 1726–1731. https://doi.org/10.1073/pnas.0610561104

68. Hindy, N. C., Ng, F. Y., & Turk-Browne, N. B. (2016). Linking pattern completion in the hippocampus to predictive coding in visual cortex. Nature Neuroscience, 19(5), 665–667. https://doi.org/10.1038/nn.4284

69. Kok, P., & Turk-Browne, N. B. (2018). Associative prediction of visual shape in the hippocampus. Journal of Neuroscience, 38(31), 6888–6899. https://doi.org/10.1523/JNEUROSCI.0163-18.2018

70. Levy, R. (2008). Expectation-based syntactic comprehension. Cognition, 106(3), 1126–1177. https://doi.org/10.1016/j.cognition.2007.05.006

71. Kleinschmidt, D. F., & Jaeger, T. F. (2015). Robust speech perception: Recognize the familiar, generalize to the similar, and adapt to the novel. Psychological Review, 122(2), 148–203. https://doi.org/10.1037/a0038695

72. McKenzie, S., Frank, A. J., Kinsky, N. R., Porter, B., Rivière, P. D., & Eichenbaum, H. (2014). Hippocampal representation of related and opposing memories develop within distinct, hierarchically organized neural schemas. Neuron, 83(1), 202–215. https://doi.org/10.1016/j.neuron.2014.05.019

73. Chanales, A. J. H., Oza, A., Favila, S. E., & Kuhl, B. A. (2017). Overlap among spatial memories triggers repulsion of hippocampal representations. Current Biology, 27(15), 2307–2317.e5. https://doi.org/10.1016/j.cub.2017.06.057

74. Fuster, J. M., & Alexander, G. E. (1971). Neuron activity related to short-term memory. Science, 173(3997), 652–654. https://doi.org/10.1126/science.173.3997.652

75. Shafi, M., Zhou, Y., Quintana, J., Chow, C., Fuster, J., & Bodner, M. (2007). Variability in neuronal activity in primate cortex during working memory tasks. Neuroscience, 146(3), 1082–1108. https://doi.org/10.1016/j.neuroscience.2006.12.072

76. Lundqvist, M., Rose, J., Herman, P., Brincat, S. L., Buschman, T. J., & Miller, E. K. (2016). Gamma and beta bursts underlie working memory. Neuron, 90(1), 152–164. https://doi.org/10.1016/j.neuron.2016.02.028

77. Miller, E. K., Lundqvist, M., & Bastos, A. M. (2018). Working Memory 2.0. Neuron, 100(2), 463–475. https://doi.org/10.1016/j.neuron.2018.09.023

